# NRG1-mediated recognition of HopQ1 reveals a link between PAMP- and Effector-triggered Immunity

**DOI:** 10.1101/293050

**Authors:** Cyril Brendolise, Marcela Martinez-Sanchez, Arry Morel, Ronan Chen, Romain Dinis, Simon Deroles, Nemo Peeters, Erik H. A. Rikkerink, Mirco Montefiori

## Abstract

- Members of the hopQ1/XopQ family of effectors are conserved amongst pathogenic bacteria suggesting an important function in virulence. Therefore, the identification of R proteins recognising members of this family is potentially of high value to achieve broad-spectrum resistance in crops.
- Recent studies demonstrated that HopQ1/XopQ recognition is EDS1-dependant and is mediated by the TIR-NB-LRR protein Roq1. Using our recently described R gene RNAi library, we have investigated the mechanism of HopQ1 recognition and the other components of its signalling network.
- We show that silencing of the CC_R_-NB-LRR helper protein N Requirement Gene 1 (NRG1) prevents HopQ1 recognition in *N. benthamiana* and that NRG1 expression confers recognition of HopQ1 and restricts *Pto* DC3000 proliferation in the otherwise susceptible host *Arabidopsis*. Moreover NRG1 expression is briefly but strongly induced at a very early stage of the plant innate response, establishing a strong dependency link between PAMP-triggered immunity (PTI) and effector-triggered immunity (ETI).
- Overall we demonstrate that in addition to Roq1, HopQ1 recognition in *N. benthamiana* requires the CC_R_-NB-LRR protein NRG1 and propose a model based on the dual requirement of a CC_R_-NB-LRR and a TIR-NB-LRR that could extend beyond HopQ1 detection and possibly be used to recognize other pathogen effectors.

## Introduction

In the course of their co-evolution, plants and pathogens have acquired increasingly complex mechanisms of resistance and susceptibility to overcome one another. In addition to physical barriers to prevent non-specific microbial attacks, plants possess two main layers of defence to counter local infections by adapted pathogens. The first layer of defence is activated upon the detection at the plasma membrane of a wide variety of evolutionarily conserved pathogen elicitors, referred to as Pathogen-Associated Molecular Pattern (PAMPs), by specific extracellular receptors (Pattern Recognition Receptors or PRRs). The PAMP-triggered immunity (PTI) leads via a complex signalling cascade to a range of cellular defence responses including Reactive Oxygen Species production, ion transport, or callose deposition. However many gram-negative pathogenic bacteria possess a type III secretion system (T3SS) which allows them to inject virulence factors (effectors) into plant cells specifically to overcome PTI. In turn, plants have developed a second layer of immunity involving intracellular receptors (resistance or R proteins) able directly or indirectly to sense the presence of bacterial effectors. The resulting effector-triggered immunity (ETI) typically leads to a rapid localized programed cell death (PCD), also referred to as Hypersensitive Response (HR), to prevent further progress of the pathogen. The co-evolution of microbes and their host plants (illustrated in the zig-zag model of Jones and Dangl, 2006) stimulates pathogens to constantly modify their repertoire of effectors to evade recognition and gain a fitness edge, and plants to increase their R protein diversity to regain resistance, leading to increasingly complex mechanisms of resistance and susceptibility.

Most of the R proteins identified to date are Nucleotide-Binding Leucine-Rich Repeat-containing proteins (NB-LRR) and usually include either a Coiled-Coil (CC) or a Toll Interleukin 1 receptor (TIR) domain in their N-termini. Activation of the PCD by NB-LRR is dependent on the well-conserved protein SUPPRESSOR OF G2 ALLELE OF SKP1 (SGT1) (Shirasu, 2009), which forms a complex with the HEAT SHOCK PROTEIN 90 (HSP90) and REQUIRED FOR MLA12 RESISTANCE 1 (RAR1) proteins to mediate stability of NB-LRR proteins (Lu *et al.*, 2003; Boter *et al.*, 2007; Zhang *et al.*, 2010). Other components of the ETI signalling pathway have been identified, such as ENHANCED DISEASE SUSCEPTIBILITY 1 (EDS1) or NONRACE-SPECIFIC DISEASE RESISTANCE 1 (NDR1), specifically required for the TIR-NB-LRR (abbreviated to TNL thereafter) and most of the CC-NB-LRR (abbreviated to CNL thereafter)-mediated ETI, respectively (Century *et al.*, 1995; Parker *et al.*, 1996; Aarts *et al.*, 1998; Zhu *et al.*, 2011; Wagner *et al.*, 2013).

Phytopathogenic bacterial strains are often defined by their host specificity, which is largely shaped by their repertoire of effectors. Host resistance specified by major resistance genes is typically conferred by direct or indirect recognition of an effector and is sometimes easily overcome by loss of that effector. It has been suggested that resistance derived from non-host species comprises various mechanism, therefore leading to a more complex and durable resistance (Mysore & Ryu, 2004; Gill *et al.*, 2015). Consequently there has been a growing interest in recent years in identifying components of resistance to pathogens from non-host species as they are believed to provide potentially more durable R genes than those originating from host species (Lee *et al.*, 2016). However, the distinction between host and non-host resistance may not be as substantial as first proposed and led Schulze-Lefert and Panstruga (2011) to propose that they differ mainly in the relative contributions from NB-LRR and PRR recognition components.

*Pseudomonas syringae pv. tomato* (*Pto*) DC3000 causes bacterial speck disease in tomato. It can also infect the model plant Arabidopsis and induces an HR in *Nicotiana benthamiana* (*Nb*). A recent study demonstrated that deletion of a single effector of its repertoire, HopQ1, is enough for *Pto* to cause disease in *Nb* (Wei *et al.*, 2007). HopQ1, like its homolog in *Xanthomonas campestris* pv. *vesicatoria* (*Xcv*) XopQ, is the sole effector, among the repertoire of these bacteria to be recognized in both *Nb* and *Nicotiana tabacum* (*Nt*) (Wei *et al.*, 2007; Adlung *et al.*, 2016). XopQ recognition within the *Solanaceae* family is largely restricted to the *Nicotiana* genus. HopQ1 and XopQ share significant homology and are functionally interchangeable. Since their recognition in *Nb* is EDS1-dependent (Adlung *et al.*, 2016; Adlung & Bonas, 2017), both effectors are likely to be recognized by a conserved mechanism within the *Nicotiana* genus involving a TNL R gene. Indeed the TNL protein Roq1 was recently identified as required to mediate HopQ1/XopQ recognition in *Nb* (Schultink *et al.*, 2017). Given the broad distribution of hopQ1/XopQ homologs within bacterial phytopathogens, including *Pseudomonas*, *Xanthomonas* and *Ralstonia* (RipB), the identification of R proteins able to recognise these effectors is potentially valuable to achieve resistance to a wide range of plant pathogens.

Previously we reported a novel approach to identify R genes involved in the recognition of effectors of interest triggering ETI in *Nb* (Brendolise *et al.*, 2017). Using this approach we demonstrate that HopQ1-triggered HR is not only dependent on Roq1 but also requires NRG1, a CNL R gene previously implicated in *Tobacco Mosaic Virus* (TMV) resistance in *Nt* (Peart *et al.*, 2005). Here, we show that silencing of NRG1 promotes *Pto* proliferation in *Nb* and that NRG1 expression confers recognition of HopQ1 and can restrict *Pto* proliferation in the susceptible host Arabidopsis. We also show that the expression of NRG1 is induced at the level of mRNA in response to PTI and discuss how this leads to an interesting new connection between PTI and HopQ1-mediated ETI that reveals a previously undetected degree of coordination between these two layers of defence response.

## Materials and Methods

### Plant material and transformation

*N. benthamiana* plants were grown in glasshouse at 22 to 24°C with natural light combined with supplemental lighting to extend the day length to 16 h. Arabidopsis plants (Columbia-0) were grown at 25°C in growth cabinets with 10 h photoperiod. Plants were transformed following the previously described floral dip method (Clough & Bent, 1998). Seeds were harvested and sterilized using a solution of 70% ethanol/0.05% triton and grown on ½ MS selecting media containing 50 mg/L kanamycin.

### Cloning and constructs

The *HOPQ1* coding sequence was amplified from *Pto* DC3000 using primers FW 5’-CACCATGCATCGTCCTATCACCGCAG-3’ and RV 5’-TCAATCTGGGGCTACCGTCGACTG-3’ and inserted in the pENTR/SD/D-TOPO vector (Invitrogen) following the manufacturer’s protocols to generate the pENTR/SD/D-HopQ1 entry clone, which was recombined by Gateway LR reaction into the pHEX2 binary vector (Hellens *et al.*, 2005). Similarly, the pENTR-GUS entry clone (Invitrogen) was recombined by Gateway LR reaction into pHEX2 to generate the pHEX2-GUS control. Hp#26, hp#27 and pTKO2-u135 have been described previously (Brendolise *et al.*, 2017). All the other individual fragments from hp#26 and hp#27 were cloned following the procedure described for pTKO2-u135, consisting of an In-Fusion HD cloning reaction (Clontech) into the pENTR/SD/D-TOPO vector to generate an entry clone, followed by an LR Gateway reaction into the pTKO2 destination vector (Brendolise *et al.*, 2017). The TRV1 and the Gateway-enabled TRV2 (pYL279) used for gene silencing in *N. benthamiana* were previously described (Liu *et al.*, 2002). The pENTR/SD/D-u135 entry clone used to generate the pTKO2-u135 construct was recombined by LR reaction into pYL279 to generate the TRV2-u135 construct.

### Transient expression and RNAi library screen

Transient expression and the RNAi library screen in *N. benthamiana* were described previously (Brendolise *et al.*, 2017). In brief, RNAi hairpin constructs, or pTKO2 empty vector, were agro-infiltrated (OD_600_ of 0.2) in leaves of 3-weeks old plants and the infiltrated area was marked. The HopQ1 construct or the GUS control were agro-infiltrated 48h later in the pre-infiltrated area. Photographs were taken 4 days after HopQ1 infiltration. For biolistic transient expression, 5 mg of gold particles (1.0 μm diameter, BioRad) were combined with 5 µg of GFP plasmid DNA pRT99-GFP (Siemering *et al.*, 1996) and 20 µg of either test DNA (e.g., HopQ1) or control DNA (pRT99-GUS) (Topfer *et al.*, 1988), treated with 20 µL of 0.1 M Spermidine solution followed by 50 µL of 2.5 M CaCl2 and placed on ice. Excess supernatant was removed to a final volume of 35 µL. Bombardment was performed on well-expanded leaves of *N. benthamiana* and Arabidopsis using a particle inflow gun made by KiwiScientific (Levin, NZ) based on the design of Finer and colleagues (Finer *et al.*, 1992). For each bombardment six target leaves were shot on the abaxial side under a metal mesh screen (100 μm) using 5 µL of the particle preparation and the following conditions: carrier to target shelf distance 14 cm (Arabidopsis) or 12 cm (*N. benthamiana*), helium pressure 300 Kpa, pulse time 30 ms, chamber vacuum −90 Kpa. Leaves were incubated in the light at room temperature for 24 h or 48 h and the number of GFP cells was counted with a fluorescence stereomicroscope (Leica Microsystems M205FA) equipped with a GFP filter set.

### VIGS

TRV2-u135 was transformed into GV3101 *Agrobacterium tumefaciens.* Bacteria were grown for two days on LB plates with appropriate antibiotics then resuspended in infiltration buffer (10 mM MgCl_2_, 10 mM MES, 150 μM acetosyringone). Strains containing TRV2-u135 or TRV2 empty vector were mixed with the strain containing TRV1 (1:1 ratio to a final OD_600_ of 0.5 each and kept for 2 h at room temperature in the dark. Two-week-old *N. benthamiana* plants were then infiltrated in one leaf with a 1 mL needleless syringe. Plants were grown for 3 weeks in a culture chamber before use.

### Plant inoculation and Bacterial growth quantification

*N. benthamiana* leaves were infiltrated with a bacterial suspension of *Pto* DC3000 ranging from 5x10^5^ cfu/mLto 10^7^ cfu/mL as indicated. The best visual difference of disease lesions between NRG1-silenced and control plants were obtained with 5x10^6^ cfu/mL and 10^7^ cfu/mL inoculum concentrations (Supporting Information Fig. S1). Immediately after infiltration and 72 hours after, 6 leaf discs of 7 mm diameter were collected from five independent patches inoculated at 5x10^6^ cfu/mL, and pulverized with glass beads using a mill grinder and O. resuspended in water. Dilutions 10^−3^ and 10^−5^ were then plated with an easySpiral^®^ instrument (Interscience) and counted according to the manufacturer recommendations. For each Arabidopsis transgenic line, six leaf discs of 6 mm diameter were collected and surface sterilized using a 0.9% bleach solution. Leaf discs were inoculated by flooding for 3min in a bacterial suspension of *Pto* DC3000 at 10^5^ cfu/mL supplemented with 0.025% Silwet L-77. Bacterial suspension was drained and leaf discs were maintained for 3 days on solid ½ MS media. Discs were surface sterilized and pulverized with glass beads in 200 µL MgCl2 buffer using the TissueLyser II system (QIAGEN). Serial dilutions were spotted onto KB media supplemented with rifampicin 25 mg/mL and the number of colonies was evaluated after 48 h of incubation at room temperature.

### Ion leakage quantification

For each treatment, sixteen 3.5 mm discs were harvested from the agro-infiltrated area of four independent *N. benthamiana* leaves and washed in distilled water for 2h. Four discs were placed in 2mL of water and conductivity was measured over time using a Laquatwin conductivity meter (Horiba). The standard errors of the means were calculated from four replicates.

### qPCR

RNA was extracted from Arabidopsis *or N. benthamiana* leaf tissue using the Spectrum Plant Total RNA kit (Sigma–Aldrich) according to the manufacturer’s instructions. The RNA was quantified using a Nanodrop ND-1000 spectrophotometer (Nanodrop Technologies, USA) and quality was evaluated on a 1% agarose gel. DNase treatment and first-strand cDNA synthesis were carried out using the QuantiTect Reverse Transcription Kit (QIAGEN) following the manufacturer’s instructions. qRT-PCR amplification was carried out on a LightCycler^®^ 480 Real-Time PCR System (Roche) using the LightCycler 480 SYBR Green I Master Mix (Roche) and analysed with the LightCycler 480 software version 1.5. Amplification efficiency of each gene was calculated from the amplification of serial dilutions of the cDNA of the corresponding gene. For the VIGS experiments, RNA was isolated from six leaf discs (7mm diam.) samples using NucleoSpin^®^ RNA kit (Macherey Nagel) and the DNAse treatment was performed using the TURBO DNA-free kit (Invitrogen). Real time qPCR amplification was performed with Transcriptor Reverse Transcriptase (Roche) with random hexanucleotides primers on 2 µg RNA. qRT-PCR was done on a LightCycler 480 using SYBR Green I Master Mix (Roche). Sequences of all the primers are provided in Table S1.

### Data Availability

All data generated in this study are included in this article (and its supplemental information file). Material is available upon reasonable request.

## Results

### NRG1 is required for HopQ1 recognition in *N. benthamiana*

*Pto* DC3000 triggers an HR in *Nb* and it was shown that HopQ1 is the sole *Pto* DC3000 effector recognised in *Nb* (Wei et al., 2007). We developed a new approach to identify R genes involved in the recognition of an effector of interest triggering an HR in *Nb* (Brendolise *et al.*, 2017). This approach is based on the genome-wide identification of most of the *Nb* R genes and their systematic silencing using an R gene RNAi library. HopQ1 was used to screen the 47 constructs of our R gene RNAi library following the methodology previously described. Two hairpin constructs, hp#26 and hp#27, were identified as being able to reduce HopQ1-triggered HR significantly (Fig. 1a). To reduce the total number of constructs in the library, hairpins were initially designed to produce multi-gene silencing by concatenation of several fragments each targeting independent genes. Each of the component fragments of hairpin constructs hp#26 and hp#27 were sub-cloned individually in the hairpin vector and their ability to reduce the HopQ1-triggered HR was tested following the same procedure. Fragments u135 and u111 from hairpins hp#26 and hp#27 respectively were identified as the fragments responsible for the reduction of HopQ1-triggered HR (Fig. 1b, c). Fragments u135 and u111 target the NRG1 and NRG2 genes respectively, with 100% identity to their targets. NRG1 is a CNL R gene previously shown to be required for TMV resistance in *Nb* (Peart *et al.*, 2005). NRG2 shares 86% homology with NRG1 and was previously shown to be a pseudogene truncated in its 5’ terminal end. Consequently the effect observed with hp#27 is likely to be due to the silencing of NRG1, given that u111 shares 92.5% identity to NRG1 over the 120 bp of the fragment (Fig. 1d). This was confirmed by the measurement of the NRG1 expression in *Nb* leaves three days after infiltration of the u111 and u135 hairpins. Both hairpins were able to reduce NRG1 expression significantly (Fig. 1e). Interestingly NRG2 transcript was also detected, however the minor decrease in NRG2 measured in the presence of the two hairpins was not statistically significant (Fig. 1e).

**Figure 1.**
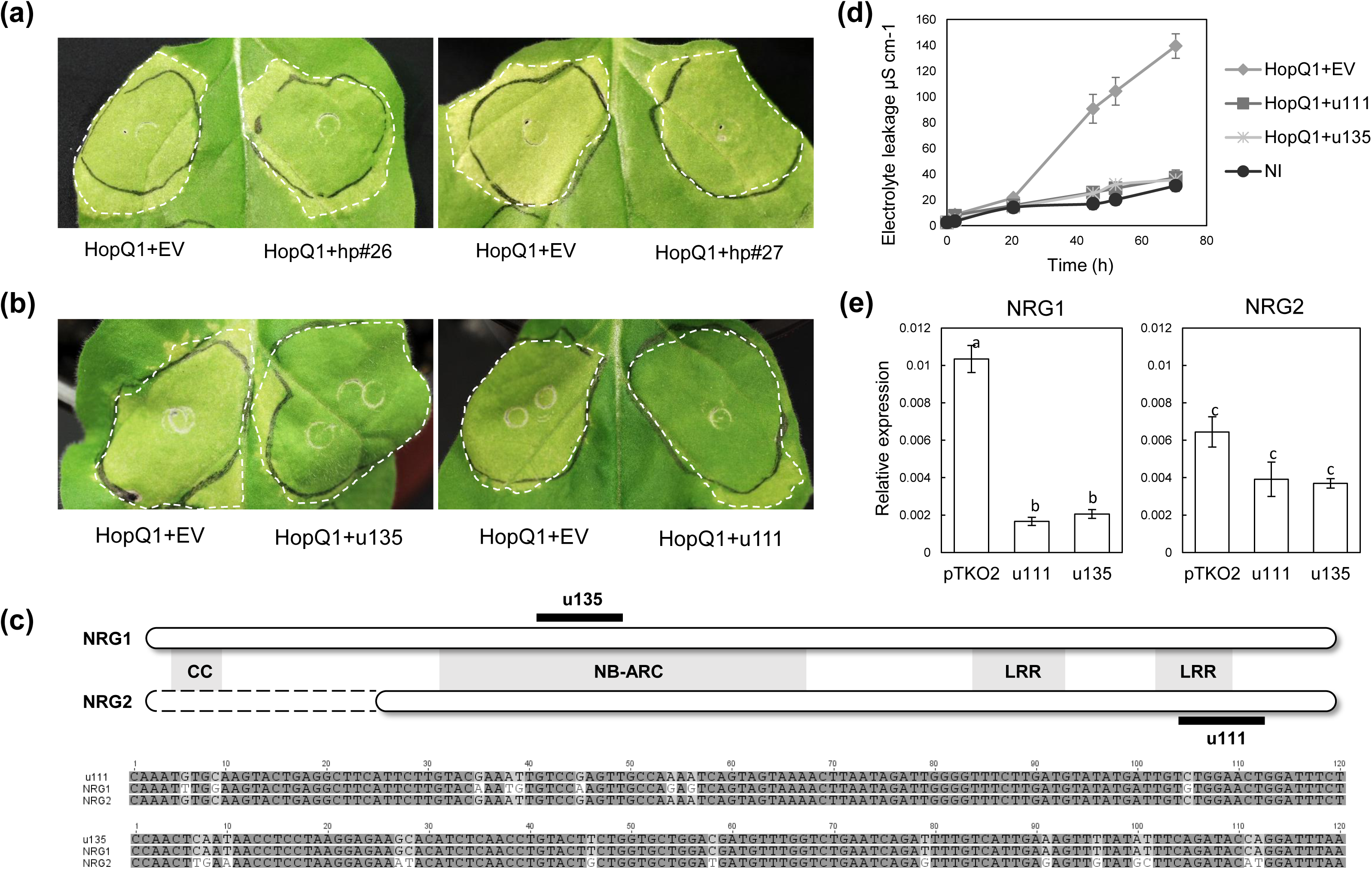
NRG1 expression is required for HopQ1 recognition. (**a**-**b**) *Nicotiana benthamiana* leaves were infiltrated by hp#26, hp#27, u135, u111 or pTKO2 empty vector (EV), and HopQ1 was infiltrated 48 h later. HR was monitored 4 days after HopQ1 infiltration. (**c**) Leakage of electrolytes was measured on the corresponding patches as indicated. The initial time point was measured two days after HopQ1 infiltration. Error bars represent the standard errors of the means calculated from four biological replicates. (**d**) Schematics of the NRG1 and NRG2 protein structure indicating the coiled-coil (CC), nucleotide binding site, presence in APAF-1, R proteins and CED-4 (NB-ARC) and leucine rich repeat (LRR) domains. Position of the u135 and u111 hairpin fragments are indicated (black bars). CLUSTALW alignment of u111 and u135 fragments with NRG1 and NRG2 nucleotide sequences. (**e**) Expression analysis of *NRG1* and *NRG2* genes in u135, u111 and pTKO2 infiltrated patches relative to the *NbPP2A* gene, 4 days after infiltration. Error bars represent the standard errors of the means calculated from four biological replicates. Values with different letters are statistically different (p<0.01) using one-way ANOVA with *post hoc* Tukey’s HSD.

### NRG1 silencing promotes *Pto* DC3000 proliferation in *N. benthamiana*

Previous studies demonstrated that deletion of HopQ1 allows *Pto* DC3000 to grow to a higher titre and cause disease in *Nb* (Wei *et al.*, 2007). To confirm the requirement of NRG1 in HopQ1 recognition, we investigated the effect of NRG1 silencing on the ability of *Pto* DC3000 to grow in *Nb*. *Agrobacterium* pre-infiltration in *Nb* is known to inhibit the *Pseudomonas* T3SS and its ability to cause disease (Chakravarthy *et al.*, 2010). Consequently we were not able to use *Agrobacterium*-mediated expression of the u135 hairpin construct to assess silencing of NRG1. Instead NRG1 was silenced by Virus Induced Gene Silencing (VIGS) using the same fragment as in the u135 hairpin construct. Two week-old *Nb* seedlings were infiltrated with the TRV1 and TRV2-u135 constructs (or the corresponding TRV2 empty vector), after 2 weeks *Pto* DC3000 was inoculated in newly formed leaves and symptoms and bacterial growth were measured. Stronger disease lesions were visible in NRG1-silenced plants than in TRV2 control plants (Fig. 2a and Supporting Information Fig. S1); moreover, *Pto* DC3000 growth measured in these plants was significantly higher than in the control plants (Fig. 2b). qPCR analysis of the corresponding infiltrated patches confirmed that NRG1 expression was reduced in plants expressing the TRV2-u135 construct (Fig. 2c).

**Figure 2.**
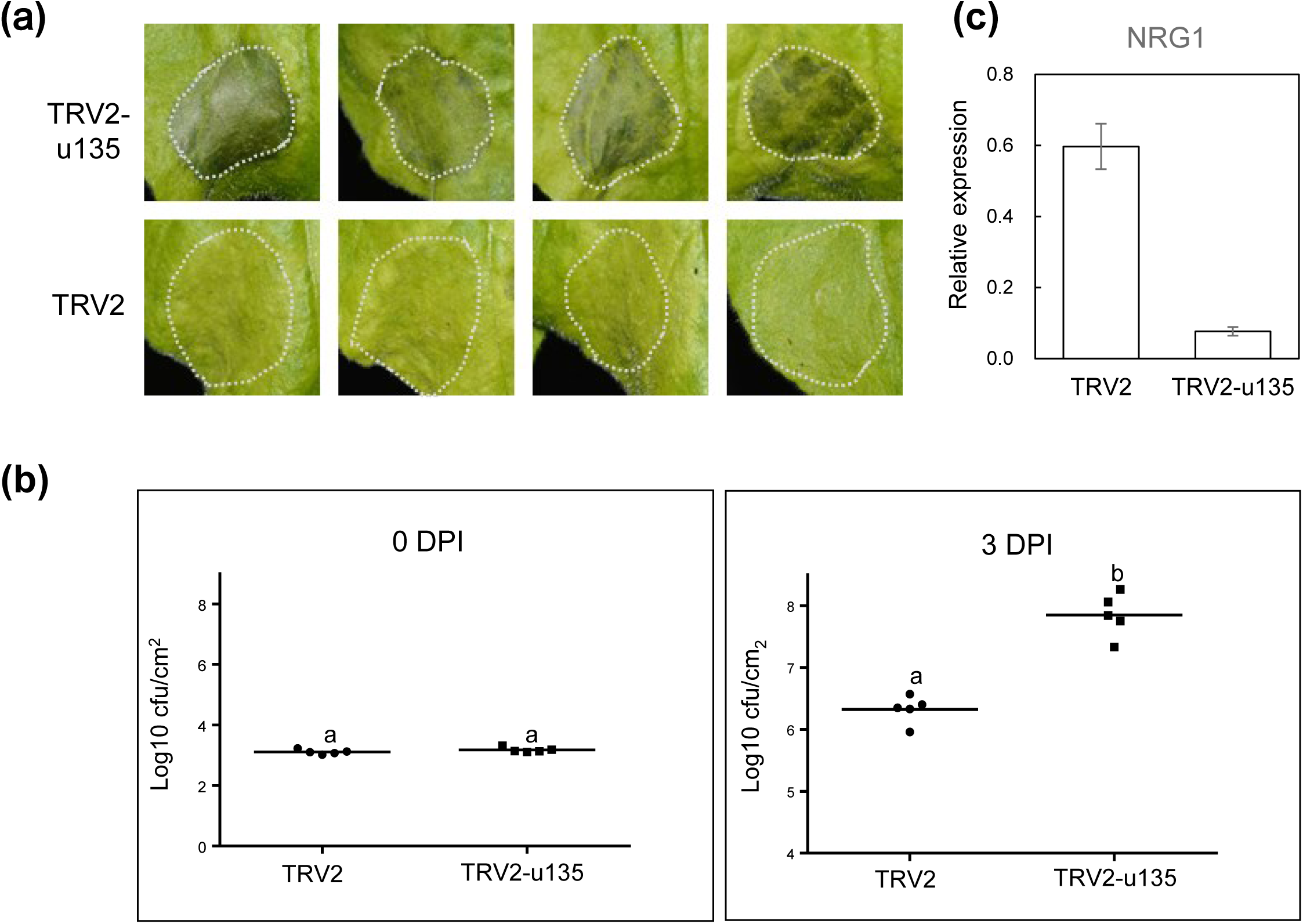
Silencing of NRG1 promotes *Pto* DC3000 growth in *Nicotiana benthamiana*. (**a**) Plants silenced for NRG1 (TRV2-u135) or infiltrated with the corresponding empty TRV2 vector were inoculated with a suspension of *Pto* DC3000 at 5x10^6^ cfu/mL. Leaves were photographed 2 days post inoculation (DPI). See also Supporting Information Fig. S1. (**b**) *Pto* DC3000 growth was evaluated immediately after inoculation and 72 h later from five leaf samples. Mean colony counts harbouring a different letter are significantly different based on a paired t-test on five biological replicates infiltrated by pair with the same inoculum dose (p<0.01). (**c**) *NRG1* expression in the silenced plants, before *Pto* DC3000 inoculation, expressed as means ± SE calculated from five leaf samples and relative to the expression of the *NbHKUK* gene.

### Expression of NRG1 in Arabidopsis confers recognition to HopQ1 and restricts *Pto* DC3000 growth *in planta*

*Pto* DC3000 is pathogenic in Arabidopsis (Whalen *et al.*, 1991). Two NRG1 orthologs have been identified in Arabidopsis (AtNRG1.1, AtNRG1.2) based on sequence similarity (53 % and 55 % similarity to NbNRG1 respectively) (Collier *et al.*, 2011). However, the expression of HopQ1 in Arabidopsis does not trigger any HR (Li *et al.*, 2013a; Hann *et al.*, 2014), suggesting that AtNRG1.1 and AtNRG1.2 are not fully functional homologs of NRG1 and/or able to recognise HopQ1.

To test whether NRG1 can confer recognition of HopQ1 in Arabidopsis, several lines of stable transformants expressing a *35S:NbNRG1* construct were generated. HopQ1 together with a GFP reporter were transiently expressed in these lines using biolistic delivery and the resulting fluorescence was quantified. HR was evaluated by quantifying a reduction of the GFP signal resulting from cell death. HopQ1 expression induced a significant reduction of the GFP reporter signal in three of the five NRG1-expressing lines tested (Fig. 3a). Conversely HopQ1 expression did not induce an HR in any of the GUS control lines (Fig. 3a and Supporting Information Fig. S2). The effector HopAS1 known to trigger a strong HR in Arabidopsis Columbia-0 (Sohn *et al.*, 2012) was used as an HR positive control. As expected HopAS1 expression induced a strong reduction of the GFP signal in both the NRG1 and the GUS lines. Interestingly, NRG1 expression level in the transformant lines inversely correlated with the GFP signal measured (Fig. 3b); indeed the two lines showing a non-significant reduction of the GFP signal also had the lowest level of expression of NRG1.

**Figure 3.**
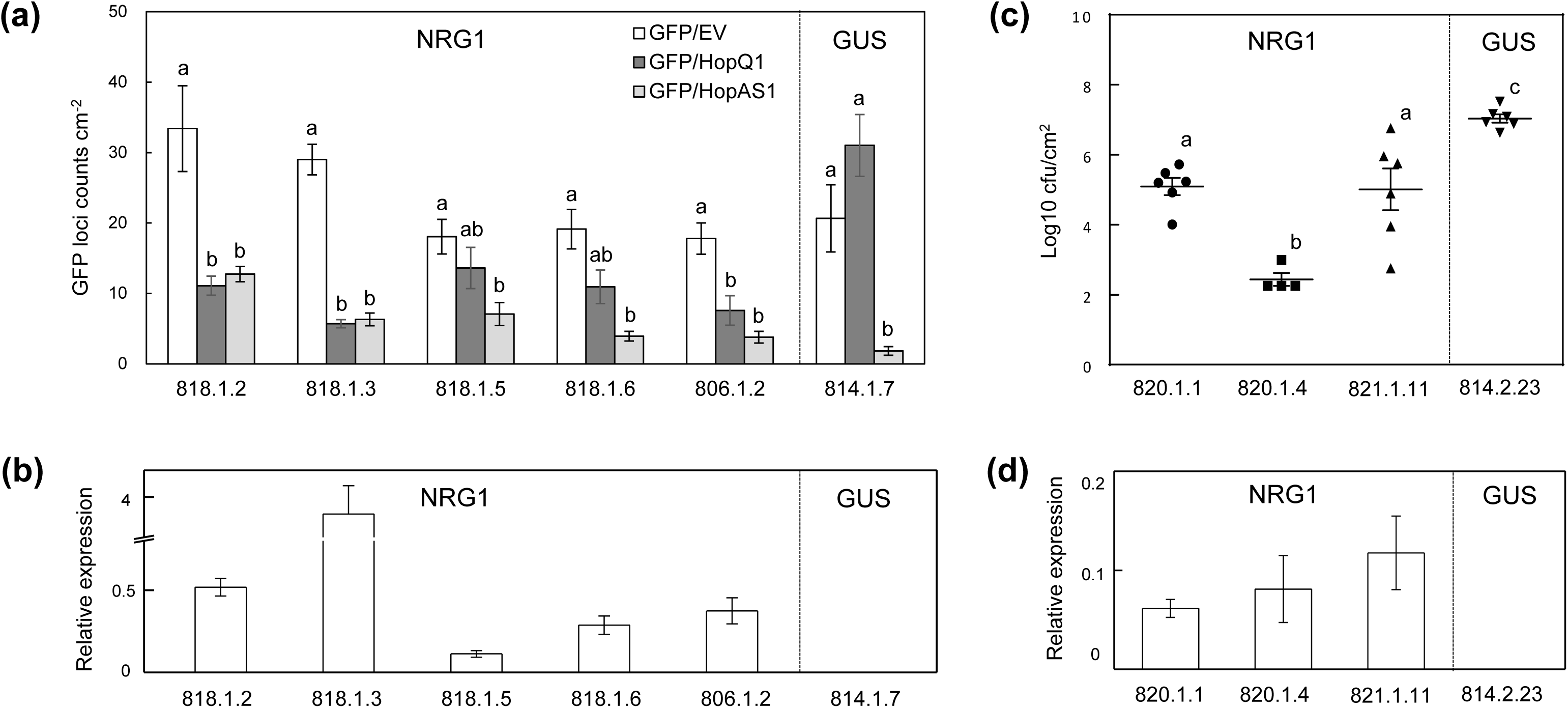
NRG1 restricts *Pto* DC3000 growth in Arabidopsis. (**a**) Effectors HopQ1 or HopAS1, or corresponding empty vector (EV), were co-expressed with a GFP reporter construct using biolistics in five independent transgenic lines of Arabidopsis expressing the 35S:NRG1 or the GUS control. The GFP signal was measured 24 h after DNA bombardment. Error bars represent the standard errors of the means calculated from six technical replicates. One-way ANOVA with *post hoc* Tukey’s HSD was applied on each individual line. Values with different letters are statistically different (p<0.01). See also Supporting Information Fig. S2. (**b**) Expression analysis of *NRG1* in the five 35S:NRG1 Arabidopsis transgenic lines and one of the GUS lines, relative to the *AtACTIN* gene, expressed as means ± SE calculated from five leaf samples. (**c**) Leaf tissues from three NRG1 lines and one GUS control were inoculated with a bacterial suspension of *Pto* DC3000 at a concentration of 1x10⁵ cfu/mL and bacterial growth was evaluated 3 days later from six leaf samples. Error bars represent the standard errors of the means and values labelled with the same letter are not statistically different based on one-way ANOVA with *post hoc* Tukey’s HSD (p< 0.01). (**d**) *NRG1* expression measured in the same NRG1 and GUS lines evaluated in the bacterial growth assay, relative to the *AtACTIN* gene and expressed as means ± SE calculated from five leaf samples.

To further assess the NRG1 effect, we tested whether NRG1 expression in Arabidopsis conferred resistance to *Pto* DC3000. Leaf discs from three NRG1 expressing lines were flood-inoculated with *Pto* DC3000 and bacterial growth was measured 4 days later. The three NRG1-expressing lines tested showed a significant decrease in bacterial count 4 days after inoculation compared with that in the GUS line (Fig. 3c). As previously, NRG1 expression in these lines was quantified and correlated inversely with the level of leaf bacterial growth (Fig. 3d). These results suggest that NRG1 expression in Arabidopsis induces recognition of HopQ1 and quantitatively restricts *Pto* DC3000 growth in the plant.

### NRG1 expression is induced at the early stages of the bacterial infection

As mentioned above, HopQ1 triggers an HR when expressed via *Agrobacterium* in *N. benthamiana*. While calibrating the biolistic delivery system for transient expression in Arabidopsis, HopQ1 and the GFP reporter were also delivered to *Nb* leaves as an HR positive control. Surprisingly, no reduction of the GFP signal was measured when HopQ1 was delivered *via* biolistics in *Nb* leaves (Fig. 4a). However, when leaves were pre-treated with *Agrobacterium* (not carrying any effector) to induce PTI prior to DNA bombardment, a significant reduction of the GFP signal was measured. These results suggest that i) unlike *Agrobacterium*-mediated transformation, the biolistic delivery of DNA bypasses the PTI and maintains the cells in a non-“primed” status, and ii) NRG1 expression, and consequently NRG1-dependent recognition of HopQ1, require activation of PTI for efficient recognition. To confirm this hypothesis, *Nb* leaves were infiltrated with Flg22 peptide to induce PTI, and NRG1 expression was analysed at various times after induction. NRG1 mRNA expression was strongly induced (> 5 fold) as soon as 30 minutes post-induction and returned to a non-induced state after 3 h, mimicking closely the expression pattern of other well-known PTI genes, ACRE 31 and CYP71D20 (Fig. 4b). By contrast a low level of NRG2 expression was detected and no induction was measured upon Flg22 treatment.

**Figure 4.**
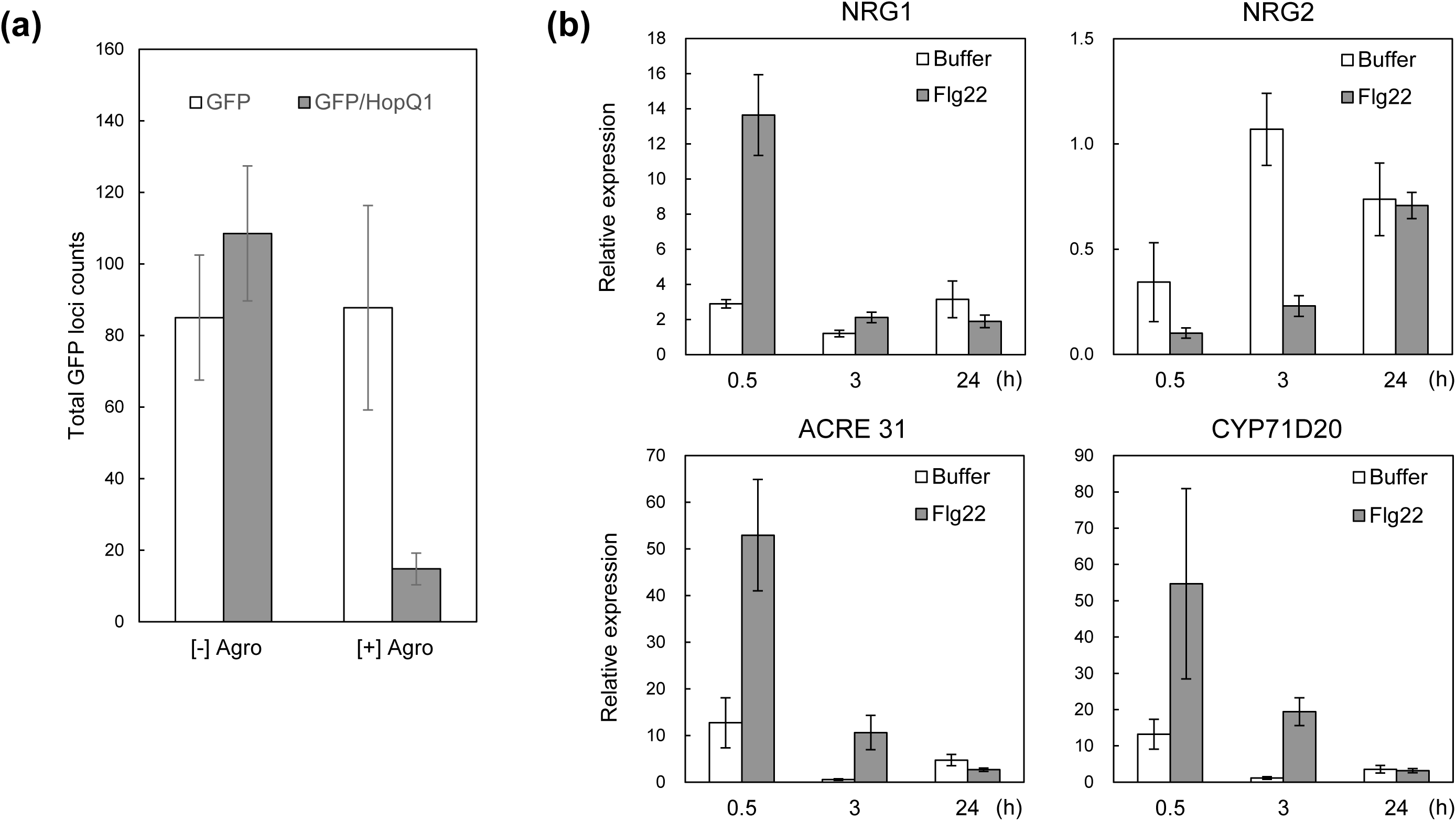
NRG1 expression is induced by the PTI. (**a**) HopQ1 and the *GFP* reporter gene were co-expressed using biolistic in *Nicotiana benthamiana* leaves pre-soaked for 10 min in an *Agrobacterium* suspension OD600 0.5 ([+] Agro) or in buffer ([-] Agro) and the GFP signal was quantified 48 h after DNA bombardment. Error bars represent the standard errors of the means calculated from four technical replicates. The experiment was repeated twice, with similar results. (**b**) *N. benthamiana* leaves were infiltrated with an Flg22 solution (10 nM) to induce the PTI and RNA were isolated at 0.5, 3 and 24 hours after induction. Gene expression levels were quantified relative to the *NbPP2A* gene and normalised for each gene to the buffer treatment at 3 hours post-induction. Error bars represent the standard errors of the means calculated from four biological replicates.

## Discussion

The HopQ1/XopQ family of effectors is highly conserved amongst plant pathogens. Recently acquired in *Pseudomonas*, it is more ancient in *Xanthomonas* or *Ralstonia* and is also found in *Acidovorax* (Rohmer *et al.*, 2004; Adlung & Bonas, 2017). The HopQ1 effector from *Pseudomonas* and its homolog XopQ from *Xanthomonas* share a high level of sequence and therefore structural similarities, suggesting a conserved yet unknown role in virulence. For instance, HopQ1/XopQ share similarities to Nucleoside Hydrolases (NH) and, although no NH enzymatic activity has yet been identified, the integrity of the NH domain is required for virulence of *P. syringae* but dispensable for HopQ1 recognition in tobacco (Li *et al.*, 2013a). HopQ1/XopQ proteins also both contain a 14-3-3 binding site, and a phosphorylation-dependent interaction with 14-3-3 proteins was demonstrated for both effectors (Giska *et al.*, 2013; Li *et al.*, 2013b; Teper *et al.*, 2014). Again, this interaction is required for virulence but not for recognition in tobacco. Finally, recognition of both effectors in *Nb* is SGT1-, EDS1- and Roq1-dependent (Wei *et al.*, 2007; Adlung *et al.*, 2016; Schultink *et al.*, 2017). The multiple overlaps in structure, mechanism and function of hopQ1/XopQ proteins suggests that their recognition in *Nb* relies on the same detection mechanism (Adlung & Bonas, 2017). Here we have demonstrated, using our previously described R gene RNAi library, that NRG1 is required for the recognition of HopQ1 in *Nb* (Fig. 1) and that silencing of NRG1 promotes the proliferation of *Pto* DC3000 in the plant (Fig. 2). Given its EDS1-dependency, our observation that HopQ1 recognition is not solely dependent on the TNL resistance gene Roq1 but also requires NRG1 which belongs to the CNL class of R proteins (Peart *et al.*, 2005) was unexpected. Because of this dual dependency on both Roq1 and NRG1 we can exclude that HopQ1 is recognized by two independent mechanisms involving these proteins. Since NRG1 was originally identified as a required component of resistance to TMV together with protein N, which belongs to the TNL class, our results rather suggest that NRG1 could be involved in a similar partnership with another TNL, the recently identified Roq1 protein, to mediate the recognition of HopQ1. NRG1 was previously reported to belong to an atypical subclass of CC-NB-LRR, named CC_R_-NB-LRR (noted RNL hereafter) for their homology to the CC domain of the RPW8 protein from Arabidopsis which confers resistance to powdery mildew (Xiao *et al.*, 2001). A related but separate branch of the CC_R_ clade contains the Arabidopsis activated disease resistance 1 (ADR1) protein and its homologs. NRG1 and ADR1 homologs were shown to form two strongly supported subclades (Collier *et al.*, 2011). Unlike other NB-LRR proteins which typically show a rapid evolutionary diversification (Jacob *et al.*, 2013), RNL members display a remarkably low level of sequence variation and moderate level of clade expansion (Collier *et al.*, 2011), suggesting that they perform a conserved function. Furthermore the RNL clade is phylogenetically basal, and perhaps therefore of more ancient origin than the rest of the CNL group. Members of the ADR1 family in Arabidopsis have a redundant function in innate immunity and ETI and were suggested to act as helper NB-LRRs in partnership with sensor NB-LRRs involved more specifically in effector detection (Bonardi *et al.*, 2011; Dong *et al.*, 2016), and are also unusual among CNL proteins in requiring EDS1 (Roberts *et al.*, 2013). RNL members have been reported in all plant species but, unlike ADR1 homologs which are found in monocots and dicots and even gymnosperms, NRG1 homologs have been identified only in dicots. Given that our evidence now shows a second occurrence of RNL-TNL interaction, the co-occurrence of NRG1-like and TNL proteins in most of the dicot species and their correlated absence in monocots and in some dicots such as Lamiales and *Aquilegia coerulea* (Collier *et al.*, 2011) is of particular interest. Co-occurrence of NRG1-like and TNL proteins in these species also coincides with the occurrence of the α/β hydrolase folded protein Senescence-Associated Gene 101 (SAG101), which was shown to interact directly with EDS1 in Arabidopsis and is typically required to trigger TNL-dependent ETI (Wagner *et al.*, 2013). Altogether these findings reinforce the general concept of a broader functional relationship between NRG1-like and TNL proteins previously only exemplified by the N and NRG1 pair but now also illustrated by the dual requirement of NRG1 and Roq1 for HopQ1 recognition. We note that no other hairpin constructs from the RNAi library were able to cancel HopQ1-mediated HR in *Nb* and our screen thus failed to identify Roq1 as a requirement for HopQ1 recognition. We have subsequently identified that the *ROQ1* gene was not annotated in the *Nb* genome at the time the RNAi library was constructed and consequently was not included in the final list of R genes used to build the RNAi library (Brendolise *et al.*, 2017).

We investigated whether NRG1-mediated recognition of HopQ1 could be transferred to other plant species. Despite the presence of two NRG1 orthologs, AtNRG1-1 and AtNRG1-2, HopQ1 is not recognized in Arabidopsis. Stable transgenic Arabidopsis lines overexpressing NRG1 were generated and transient expression of HopQ1 in the resulting lines was shown to trigger an HR. In addition, the growth of *Pto* DC3000 was also restricted when inoculated in these lines (Fig. 3). These results indicate that NRG1 expression in Arabidopsis can confer HopQ1 recognition and confer resistance to *Pto* DC3000. These results also suggest therefore that a functional ortholog of Roq1 may be present in Arabidopsis. Roq1 was reported to be highly conserved in the *Nicotiana* genus but not detected in other plant species (Schultink *et al.*, 2017). Several potential candidates can be identified by BLASTp search of the Arabidopsis annotated genome, but their low sequence homologies to Roq1 (highest was AT5G17680.1 with 47.1% homology) do not identify an obvious sequence ortholog despite NRG1 being able to mediate the recognition of HopQ1 in Arabidopsis. This could mean that Roq1 functional homologs are not restricted to the *Nicotiana* genus and might be more widely distributed in plants despite poor sequence homology. The ability of NRG1 to function in Arabidopsis indicates functional conservation within the NRG1 family. It would therefore be interesting to investigate whether NRG1-mediated HopQ1 recognition can function in other species, in particular those known to possess only the CNL type of R genes, such as monocots or Lamiales. Similarly, Roq1 was reported to mediate the recognition of HopQ1 when expressed in *Beta vulgaris* and *Nicotiana sylvestris* (Schultink *et al.*, 2017). This suggests that both species contain a functional homolog of NRG1. TBLASTN search of *N. sylvestris* genome identifies two sequence othologs of NbNRG1 (XP_009777016.1 and XP_009769791.1) with 96% and 76% similarities respectively. However the best sequence ortholog we identify in the more distant species *B. vulgaris* only shares 50% similarities with NbNRG1, yet seems to be functionally able to mediate, together with Roq1, the recognition of HopQ1. These results have important consequences on engineering resistance in plants by using such dual resistance components as the deployment of both partners may be required to transfer the resistance to any crop of interest.

An unusual property of NRG1 is that its expression is induced in the early stages of plant infection. We show that NRG1 is at its highest level of expression as early as 30 min after induction of PTI by Flg22 (Fig. 4) and returns to a non–induced state 3 h after induction of PTI, suggesting that NRG1 expression is subjected to a tight regulation. That relationship between mRNA expression and resistance is also evidenced by the link between transgenic expressions levels of NRG1 in Arabidopsis and the ability to recognize HopQ1. The hypothesis of tight regulation requirement is further reinforced by the finding that overexpression of NRG1, or its CC_R_ domain alone, induces an HR in *Nb* (Peart *et al.*, 2005; Collier *et al.*, 2011). Interestingly, although the function of HopQ1/XopQ in virulence remains unknown, several lines of evidence that point towards interfering with plant innate immunity have been reported. HopQ1 was reported to be one of three effectors that directly or indirectly cause the degradation of RPM1-interacting protein 4 (RIN4), a negative regulator of plant innate immunity in Arabidopsis and tomato, when co-expressed in *Nb* (Luo *et al.*, 2009). HopQ1 was also demonstrated to inhibit innate immunity by reducing FLS2 level possibly through activation of cytokinin signalling (Hann *et al.*, 2014). While several other instances of effectors targeting the innate immunity have been reported, the antagonistic interplay between the HopQ1 function in virulence and its detection mechanism in *Nb* would be of particular interest. Since Roq1 was shown to interact physically with HopQ1 (Schultink *et al.*, 2017), we propose a model where NRG1, rapidly expressed upon PTI, would together with Roq1 prime the defence response (Fig. 5a). The physical interaction of HopQ1 with Roq1 would somehow disrupt this balance and initiate the signalling cascade leading to cell death (Fig. 5b). It is interesting to note that, similarly, the direct interaction between the TMV replicase p50 and N was shown to be required to trigger the NRG1-mediated resistance to TMV (Erickson *et al.*, 1999; Ueda *et al.*, 2006). Whether the TNL partner (Roq1 or N) is constitutively expressed or also induced by PTI remains to be determined. Any configuration where either of the two partners is missing would fail to recognize HopQ1 and trigger ETI (Fig. 5c-d). Although very speculative, such a model would be compatible with the SGT1-independent nature of the HR triggered by overexpressing NRG1 or its CCR domain alone (Collier *et al.*, 2011) and also compatible with the SGT1- and EDS1-dependence of HopQ1 recognition due to the involvement of a TNL partner. Additional analysis of protein-protein interaction between the different partners will be required to support this model.

**Figure 5.**
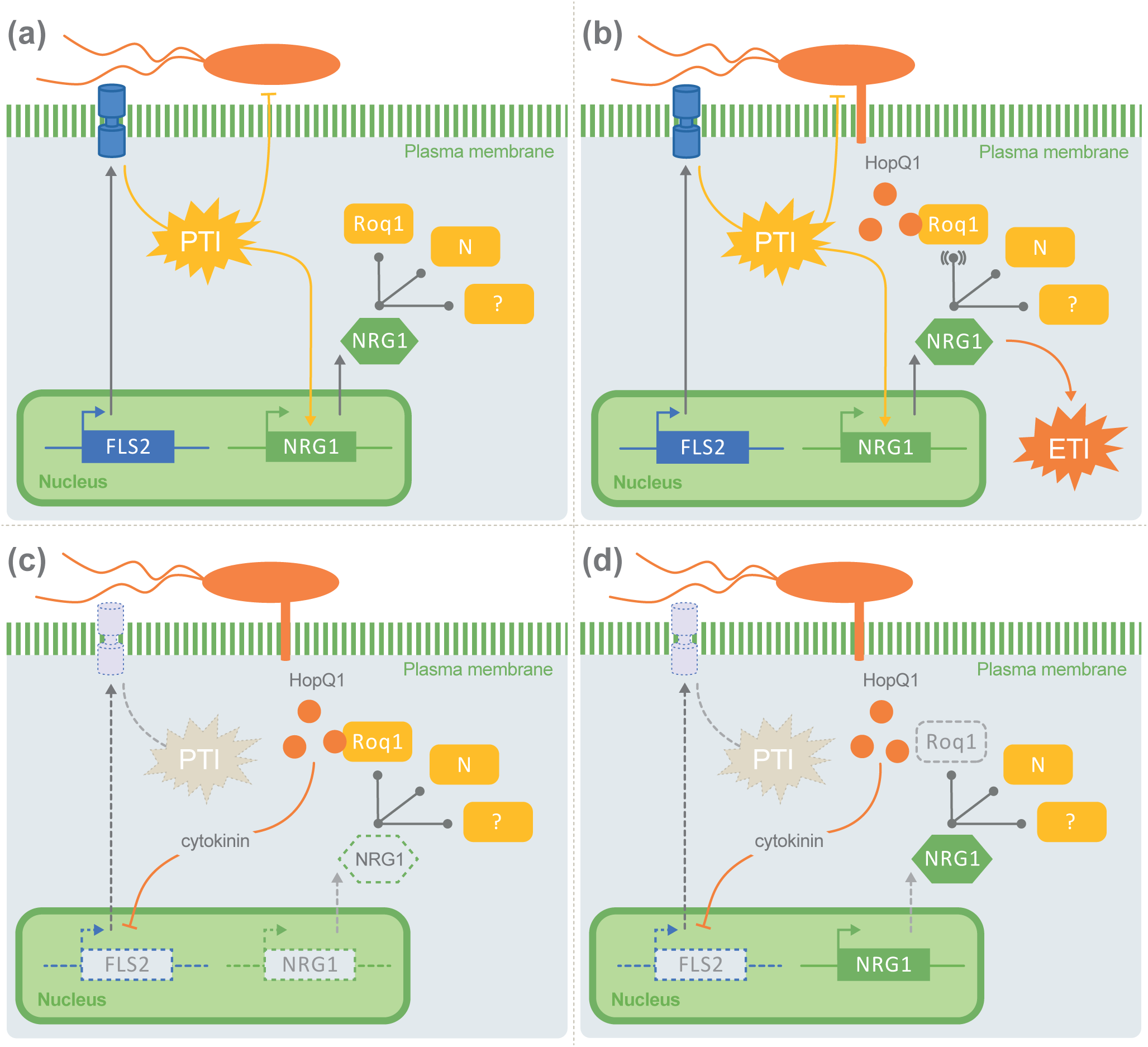
Model of HopQ1 recognition. Bacteria are sensed by PRR such as FLS2 and trigger PTI. NRG1 is rapidly expressed and primes the ETI response but remains in a non-activated state through an equilibrium with its TNL partners, such as N or Roq1 (**a**). Upon injection of type III effectors, Roq1 binds HopQ1 and disrupts the balance with NRG1 by a mechanism which remains to be determined, and triggers ETI (**b**). In a configuration where NRG1 is not present or not functional, the interaction between Roq1 and HopQ1 would occur but would not be detected by NRG1 therefore no ETI would be induced and HopQ1 would progressively dampen the PTI by repressing FLS2 expression. This configuration potentially corresponds to Arabidopsis in which NRG1 overexpression is sufficient to mediate HopQ1 recognition (**c**). Similarly in the absence of a functional Roq1, NRG1 would be induced upon PTI activation, prime the ETI response through interaction with other TNLs (such as N) but in the absence of Roq1, HopQ1 would not be detected by NRG1 and would eventually repress FLS2 expression. This configuration would be consistent with plants species such as *Beta vulgaris* of *Nicotiana sylvestris* in which Roq1 overexpression is sufficient to mediate HopQ1 recognition (**d**).

In conclusion, we have demonstrated that the recognition of HopQ1 in *Nb* is mediated by NRG1. This result together with the recent study revealing the role of the Roq1 in HopQ1 recognition, is the second illustration of the concept of dual NB-LRR requirement consisting of a TNL and a helper RNL to mediate effector recognition (Baggs *et al.*, 2017) and suggests that NRG1 could be involved in the recognition of additional effectors through similar partnerships with other TNLs. The association between the dual R protein requirement and the ability of NRG1 to be induced by PTI signals provides a novel and previously undiscovered degree of cooperation between PTI and ETI resistance.

## Acknowledgements

We thank G Wadasinghe and Monica Dragulescu for their efforts in the maintenance of plants in the glasshouse. We thank Kee H. Sohn for providing the HopAS1 construct. Finally, we thank Zac Hanley and Jibran Tahir for critically reviewing the manuscript and Darren Snaith for the visual illustrations. This work was supported by Plant and Food Research internal Core funding and the New Zealand Ministry for Business, Innovation and Employment (C11X1205). This work also was supported by the "Laboratoire d’Excellence (LABEX)" TULIP (ANR-10-LABX-41). The authors declare no competing interests.

## Author Contributions

CB and MM designed the research and wrote the manuscript together with EHAR and NP. CB and RD performed the library screen with HopQ1 and subsequent identification of NRG1. MMS performed stable transformation of Arabidopsis and characterization of the transformants. AM and NP performed the VIGS experiments. Gene expression analysis were done by MMS, MM and AM. RC and SD performed the biolistic transient expressions and analysis.

## Supporting Information

Figure S1. *Pto* DC3000 promotes disease lesion in NRG1-silenced *Nicotiana benthamiana*.

Figure S2. HopQ1 is not recognized in Arabidopsis control lines.

Table S1. Sequence of the primers used in this study.

